# Semaphorin 3a reduces the side effects of radiation on BMSCs by reducing ROS

**DOI:** 10.1101/837492

**Authors:** Bo Huang, Haiyang Tang, Tao He, Zheng Yang, Ping Gong

## Abstract

1.

**Background/Aims:** Radiotherapy does not only kill tumor cells but also impairs the function of adjacent tissues, especially bone metabolism by damaging bone marrow stromal stem cells (BMSCs). This study aimed to investigate the effect of semaphorin 3a (Sema3a) on BMSCs exposed to 2 Gy radiation.

**Materials:** BMSCs were divided into four groups, namely, group A (0 Gy), group B (2 Gy), group C (0 Gy + Sema3a), and group D (2 Gy + Sema3a). A Cell Counting Kit-8 kit, Alizarin-Red and Oil-Red-O staining, alkaline phosphatase activity kit, and dichlorodihydro-fluoresce in diacetate were used to test cell proliferation, cell cycle, osteogenic ability, adipogenic ability, and the level of reactive oxygen species (ROS), respectively, in each group. Real-time PCR was performed to test the expression of osteogenic (osteocalcin and Runt-related transcription factor 2), adipogenic (peroxisome proliferator-activated receptor gamma), interleukin (IL)-6, and tumor necrosis factor (TNF)-α genes.

**Results:** BMSC proliferation, osteogenic differentiation, and the number of cells undergoing division (S+G2 phase of the cell cycle) were found to be lower in group B than in group A. and the cellular levels of ROS, adipogenic differentiation, and expression of inflammatory factors (IL-6 and TNF-α) were higher in group B than in group A. Furthermore, osteogenic differentiation ability was higher in group D than in group B, and adipogenic differentiation ability, cellular levels of ROS, and gene expression of TNF-α and IL-6 were lower in group D than in group B.

**Conclusion:** This study demonstrated that 2 Gy radiation could decrease the osteogenic differentiation ability of BMSCs and increase their adipogenic differentiation ability by increasing the production of ROS. However, Sema3a could reduce these side effects by decreasing the levels of ROS.

## 2. Introduction

Radiotherapy, incorporating high-energy ionizing radiation, is used to treat many types of cancer. Radiotherapy can kill tumor cells, inhibit their proliferation, induce their death, and treat the pain caused by bone cancer [1–2]; however, it also damages surrounding tissues and causes systemic metabolic disorders, especially on bone metabolism [3–4]. A total dosage of 60 Gy in fractions can be used in combination with surgery to remove a primary tumor, such as osteosarcoma [5]. A dosage of 50 Gy radiation leads to an almost 50% loss of mineral content and reduction of the elastic modulus of bone [6]. Radiotherapy induces the production of oxygen-free radicals in local tissue. Reactive oxygen species (ROS), produced by the radiolysis of water, activate the transcription factor nuclear factor kappa-B, in turn enhancing the expression of p16INK4A. The expression of p16INK4A protein activates the pRb tumor suppressor protein, thereby suppressing the expression of certain genes involved in cell proliferation, ultimately leading to durable cell-cycle arrest triggering mutagenesis, DNA damage, apoptosis, and nucleotide excision repair, causing an inflammatory response and the expression of inflammatory factors [26–27, 42, 50]. These post-radiation effects can lead to osteopenia, radiation-induced osteoporosis, and a higher risk of serious fractures [7–8]. Recent reports showed that the incidence of fracture is as high as 22% in patients with breast cancer and 24% in patients with soft-tissue sarcoma [7–9]. The rate of rib fracture increases 10-fold in patients with breast cancer receiving radiotherapy in comparison to healthy individuals [9]. In addition to radiotherapy, cosmic rays and radiation from nuclear weapon can also cause abnormal bone metabolism and increase the risk of fractures [10].

The healing time for post-radiation fractures in patients with carcinoma is usually more than 6 months, and bone union is delayed in up to 67% of patients [11]. Clinical data have demonstrated that the failure rate of dental implants is two to three times higher in irradiated bone than in non-irradiated bone [12]. Although the side effects of radiation have been well studied, there is no consensus on treatment and an accurate prognosis is difficult to generate [13]. Hyperbaric oxygen (HBO) treatment is used to relieve the side effects of radiotherapy as it raises oxygen concentration, improves bone formation, and promotes the healing of soft tissue by inducing angiogenesis and increasing bone metabolism. However, HBO therapy is contraindicated in several ailments, including pulmonary disease, ocular aneurysm, convulsions associated with oxygen toxicity, and rupture of the drum membrane [14–15]. In addition, the time, cost, and real necessity for HBO therapy should also be considered. Other studies have even reported that HBO provides no additional benefits for improving the success rate of dental implants in irradiated tissues [15].

Semaphorin 3a (Sema3a), a prototype axonal guidance molecule in the semaphorin family, is expressed in a wide range of tissues, including bone, cartilage, endothelial cells, glia, teeth, neurons, connective tissue, and muscle, and plays a positive role in bone metabolism [16–18]. It is involved in many physiological processes, such as cell apoptosis, cell adhesion, cell migration and patterning, guidance of axonal growth, vascular reconstruction and growth, tumor metastasis, cytokine release, and immune cell regulation [16–21]. Hayashi et al demonstrated that Sema3a expressed in osteoblasts functions as a potent osteoprotective factor by synchronously promoting bone formation and inhibiting bone resorption in the bone formation phase [17]. When Sema3a binds to neuropilin 1, it stimulates osteogenic differentiation and inhibits adipocyte differentiation of bone marrow stromal stem cells (BMSCs) through the canonical Wnt/β-catenin signaling pathway, which functions through the pleckstrin domain-containing protein 2 (FARP2)-mediated activation of Ras-related C3 botulinum toxin substrate 1 (Rac1) during osteoblast differentiation [17]. Another study indicated that Sema3a expressed in neurons regulates bone formation and resorption indirectly by modulating sensory nerve development, but it does not act directly on osteoblasts [16]. In vivo studies have demonstrated that global knockout of Sema3a results in a reduction in the quantity and quality of bone. In wild-type mice, bone volume per tissue volume is 30%; however, in Sema3a global knockout mice, this value is only 8% [16–17]. In an ovariectomized mouse model of postmenopausal osteoporosis, Sema3a suppresses osteoclastogenesis and promotes osteoblastogenesis [22].

Sema3a plays an important role in the regulation of bone remodeling; however, it is unclear whether Sema3a can alleviate the damage to BMSCs caused by radiotherapy [23]. The present study aimed to examine the effect of Sema3a on BMSCs under 2 Gy gamma radiation.

## 3. Materials and Methods

### 3.1 Cell culture

BMSCs were isolated from Sprague-Dawley rats (weighing 120 ± 10 g) supplied by Sichuan University Animal Center. The rats were killed through cervical dislocation, tibiae and femurs were removed, and bone marrow cells were flushed out with Dulbecco’s modified Eagle’s medium (DMEM; HyClone, Logan, UT). The cells were cultured in DMEM supplemented with 10% fetal bovine serum (Gibco, Melbourne, Australia) and incubated at 37°C in an atmosphere of 5% CO2. Non-adherent cells were discarded after culture for 12 h.

### 3.2 Radiation

The cells were digested with 0.25% trypsin and re-suspended in DMEM. A single dose of 2 Gy gamma radiation was administered at a rate of 0.83 Gy/min in the Seventh People’s Hospital in Chengdu, China. The source-bottle distance was 80 cm and the field size was 10 × 10 cm2. At the same time, the samples in the control group were kept outside the radiation room under the same conditions. The cells were then divided into four groups: group A, 0 Gy radiation; group B, 2 Gy radiation; group C, 0 Gy radiation + 50 ng/mL Sema3a; and group D, 2 Gy radiation + 50 ng/mL Sema3a.

### 3.3 Cell proliferation assay

BMSCs were seeded in 96-well plates at a density of 3.0 × 103 cells/well with different concentrations of Sema3a (0, 10, 50, or 100 ng/mL). The proliferation of BMSCs was assessed on the 1st, 3rd, 5th, and 7th day using a Cell Counting Kit-8 (CCK-8) assay (Dojindo, Kumamoto, Japan). Optical density (OD) was measured at 450 nm using a microplate reader (Varioskan Flash; Thermo Fisher Scientific, Waltham, MA). Cell proliferation in the four groups was determined by the same method.

### 3.4 Osteogenic differentiation, alkaline phosphatase activity assay, and adipogenic differentiation

The four groups were seeded in 6-well plates at a density of 2.0 × 104 cells/well and cultured in osteogenic or adipogenic medium. The cells were fixed in 4% paraformaldehyde after differentiation. Osteogenic differentiation was assessed using 1% Alizarin-Red (Sigma-Aldrich, St. Louis, MO) on the 21st day, and adipogenic differentiation was determined with 0.3% Oil-Red-O (Sigma-Aldrich) on the 7th day. Images were obtained with a reverse phase contrast microscope (ZE4 HD, high definition; Leica, Wetzlar, Germany), and analyzed by Image Pro Plus 6 software (Media Cybernetics, Rockville, MD). The quantity of calcium mineral was measured by using cetylpyridinium chloride. The quantity of triglyceride (TG) in the cells was calculated using a serum TG determination kit (Sigma-Aldrich). To measure alkaline phosphatase (ALP) activity, the cells were collected on the 7th day, washed twice with cold phosphate-buffered saline (PBS), lysed by freezing-thawing and ultrasound pyrolysis three times, and measured using an ALP activity kit (Nanjing Jiancheng Research Institute, Nanjing, China). The total amount of protein was measured with a bicinchoninic acid protein measurement kit (KeyGen Biotech, Nanjing, China).

### 3.5 Cell cycle assay

At 24 h after corresponding treatments, the cells were digested with 0.25% trypsin and re-suspended in PBS (1.0 × 106 cells/mL). The cell cycle was measured using a Cell Cycle Detection Kit (KeyGen Biotech) under a fluorescent microscope (Zeiss Axioplan, Zeiss, Oberkochen, Germany; 10×40) equipped with an FITC and DAPI filter according to the manufacturer’s instructions.

### 3.6 ROS assay

At 2 h after treatment, the levels of ROS were examined. Intracellular ROS were detected by incubating the cells with the fluorescent probe dichlorodihydrofluorescein diacetate (DCFH-DA) according to the manufacturer’s instructions (Sigma-Aldrich). The cells were washed three times with cold PBS. Images were obtained with a reverse phase contrast microscope (ZE4 HD, high definition; Leica) and analyzed by using Image Pro Plus 6 software. Cell culture samples were examined under a fluorescent microscope (Zeiss Axioplan; 10×40).

### 3.7. Real-time PCR assay

On the 5th and 10th day, total RNA was extracted with an RNA extraction kit (Bioer Technology, Hangzhou, China) following the manufacturer’s protocol. RNA concentration was measured with a spectrophotometer. RNA with an OD value (A260/A280) of 1.8–2.0 was reversed transcribed to cDNA by using a PrimeScriptTM RT-PCR kit (Takara, Kusatsu, Japan). cDNA was amplified by Takara Taq™ (DR001AM; Takara) for 40 cycles (denaturation for 30 s at 95°C, primer annealing for 5 s at 95°C, and extension for 31 s at 60°C). RT-PCR was carried out in quadruplicate on an ABIPRISM 7300 Real-time PCR system (Applied Biosystems, Foster City, CA). The sequences of the primer pairs are provided in Table 1; glyceraldehyde 3-phosphate dehydrogenase (GAPDH) was used as an internal control, and Runt-related transcription factor 2 (RUNX2), osteocalcin (OCN), interleukin (IL)-6, tumor necrosis factor (TNF)-α, and peroxisome proliferator-activated receptor gamma (PPARγ) were examined.

**Table 1.**
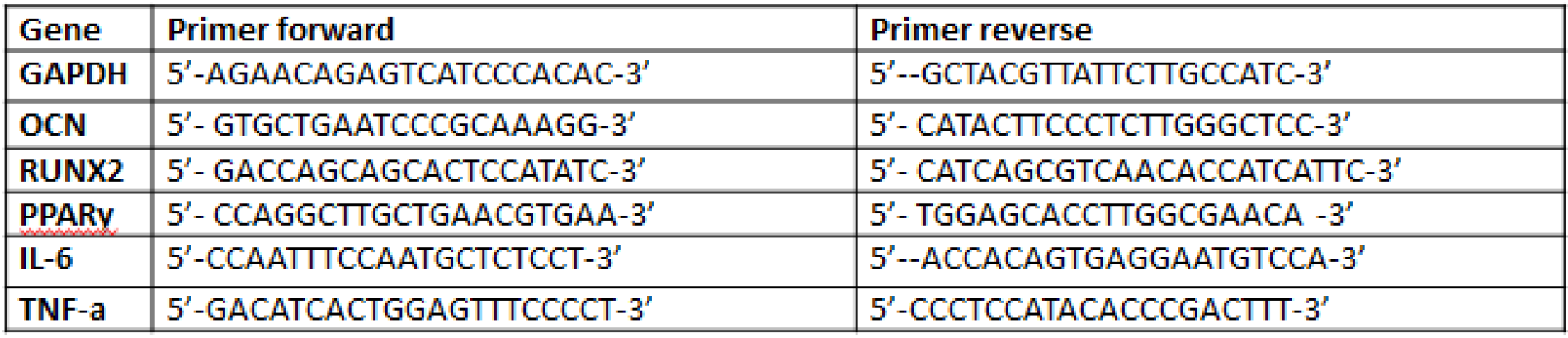
Primer pairs used in the study.

### 3.8 Statistical analysis

Data were analyzed using SPSS version 21.0 software (SPSS Inc., Chicago, IL) using one-way analysis of variation. All data are expressed as the mean ± standard error of the mean. P-values < 0.05 were considered statistically significant. Four to five independent replicates were performed for each experiment.

## 4. Results

### 4.1. Characterization of rat BMSCs

In order to verify the multiple differentiation potential of the isolated BMSCs, their ability to differentiate into osteoblasts and adipocytes was assessed by using Alizarin-Red and Oil-Red-O staining, respectively. As shown in Figure 1, positive staining was observed for osteogenic and adipogenic differentiation.

**Fig. 1.**
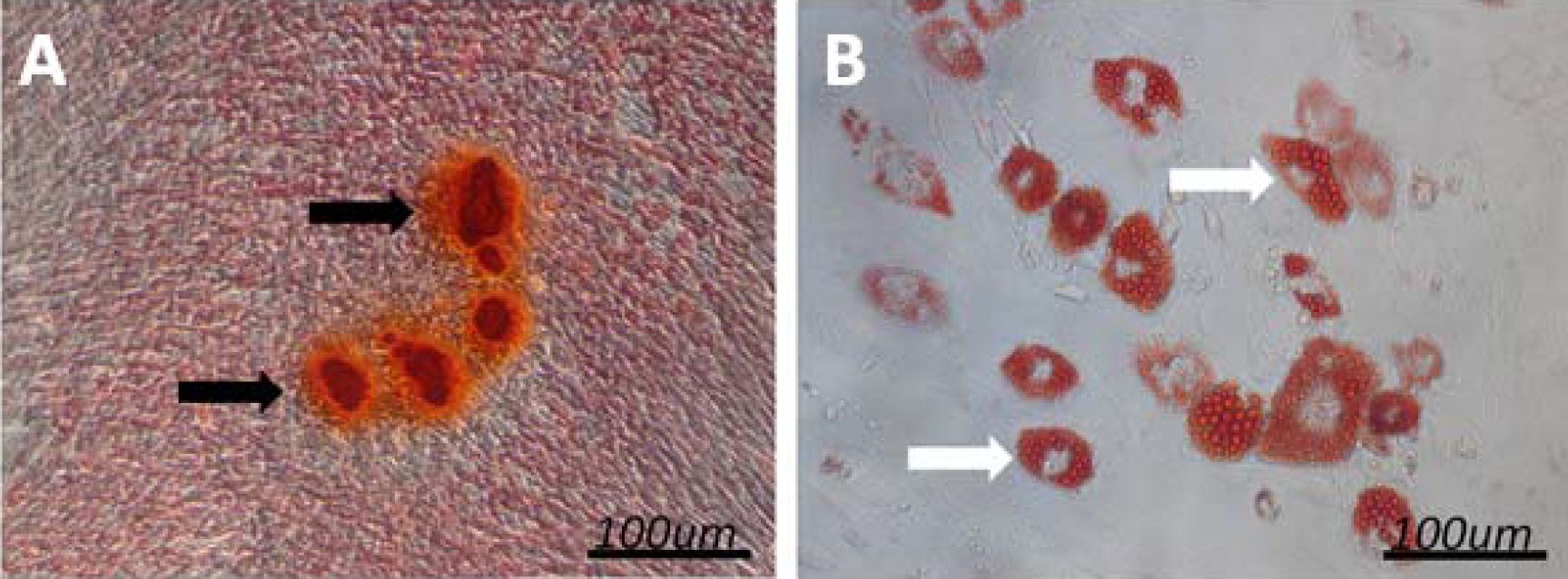
Characterization of BMSCs. (A) Alizarin-Red-S-positive BMSCs were observed after the induction of osteogenesis for 3 weeks. The black arrows indicate bony nodules. (B) Oil-Red-O-positive BMSCs were observed after the induction of adipogenesis for 10 days. The white arrows indicate lipid droplets.

### 4.2. Effect of Sema3a and 2 Gy radiation on BMSC proliferation

To test the effect of Sema3a and 2 Gy radiation on cell proliferation, we initially examined the effects of different concentrations of Sema3a and 2 Gy radiation on the proliferation of BMSCs by using a CCK-8 assay kit. As shown in Figure 2a, on the 5th and the 7th day, cell proliferation was higher in the 10 and 50 ng/mL Sema3a groups than in the 0 ng/mL group, but the difference was only statistically significant between the 50 and 0 ng/mL groups (P < 0.05). However, cell proliferation was lower in the 100 ng/mL Sema3a group than in the 0 ng/mL Sema3a group (P < 0.05). Therefore, a concentration of 50 ng/mL Sema3a was used in the subsequent experiments, and this result was similar to that of a previous study [16]. As shown in Figure 2b, cell proliferation was lower on the 5th and 7th days in group B than in group A (P < 0.05); however, it was higher in group D than in group B on both days (P < 0.05).

**Fig. 2.**
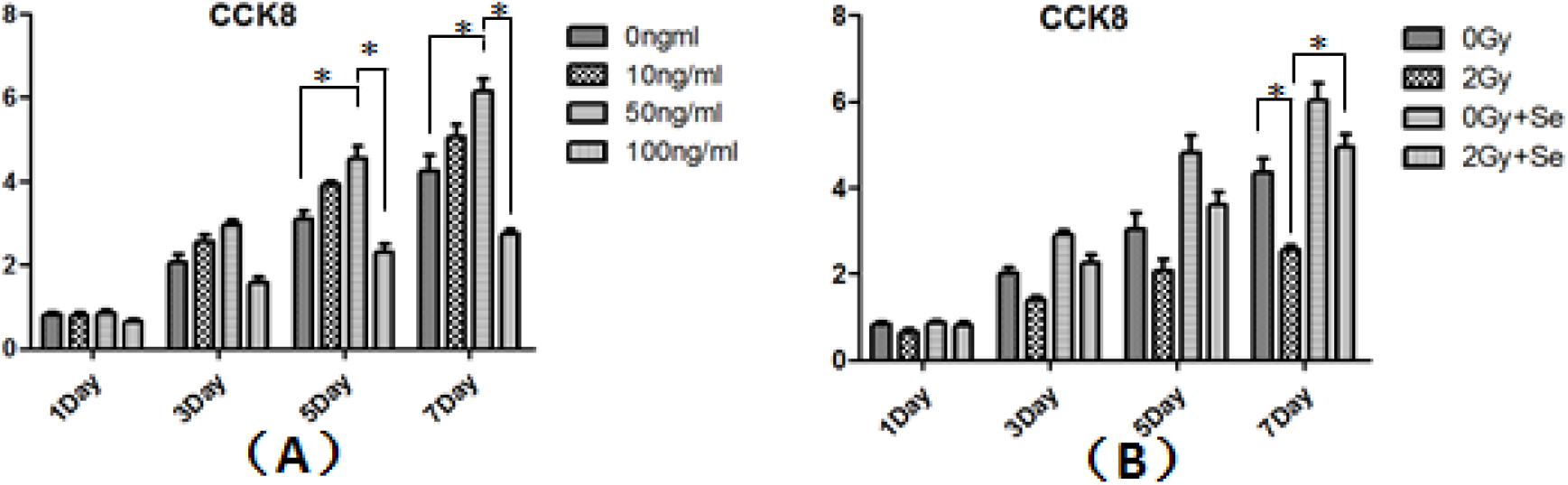
Effect of Sema3a and 2 Gy radiation on BMSC proliferation. (A) Effect of different concentrations of Sema3a on the proliferation of BMSCs. (B) Effect of 2 Gy radiation and Sema3a on the proliferation of BMSCs. *P < 0.05. All values are expressed as the mean ± standard deviation (SD), N = 5.

### 4.3 Effect of radiation and Sema3a on BMSC osteogenesis

Calcium deposition and ALP activity were used to assess the effect of Sema3a and 2 Gy radiation on the osteogenic differentiation of BMSCs. As shown in Figure 3, ALP activity and calcium deposition were reduced in group B compared with group A (P < 0.05), with the reduction in ALP activity and calcium deposition being approximately 60% and 30%, respectively. ALP activity and calcium deposition were higher in group D than in group B (P < 0.05), while ALP activity and calcium deposition were similar in groups A and D.

**Fig. 3.**
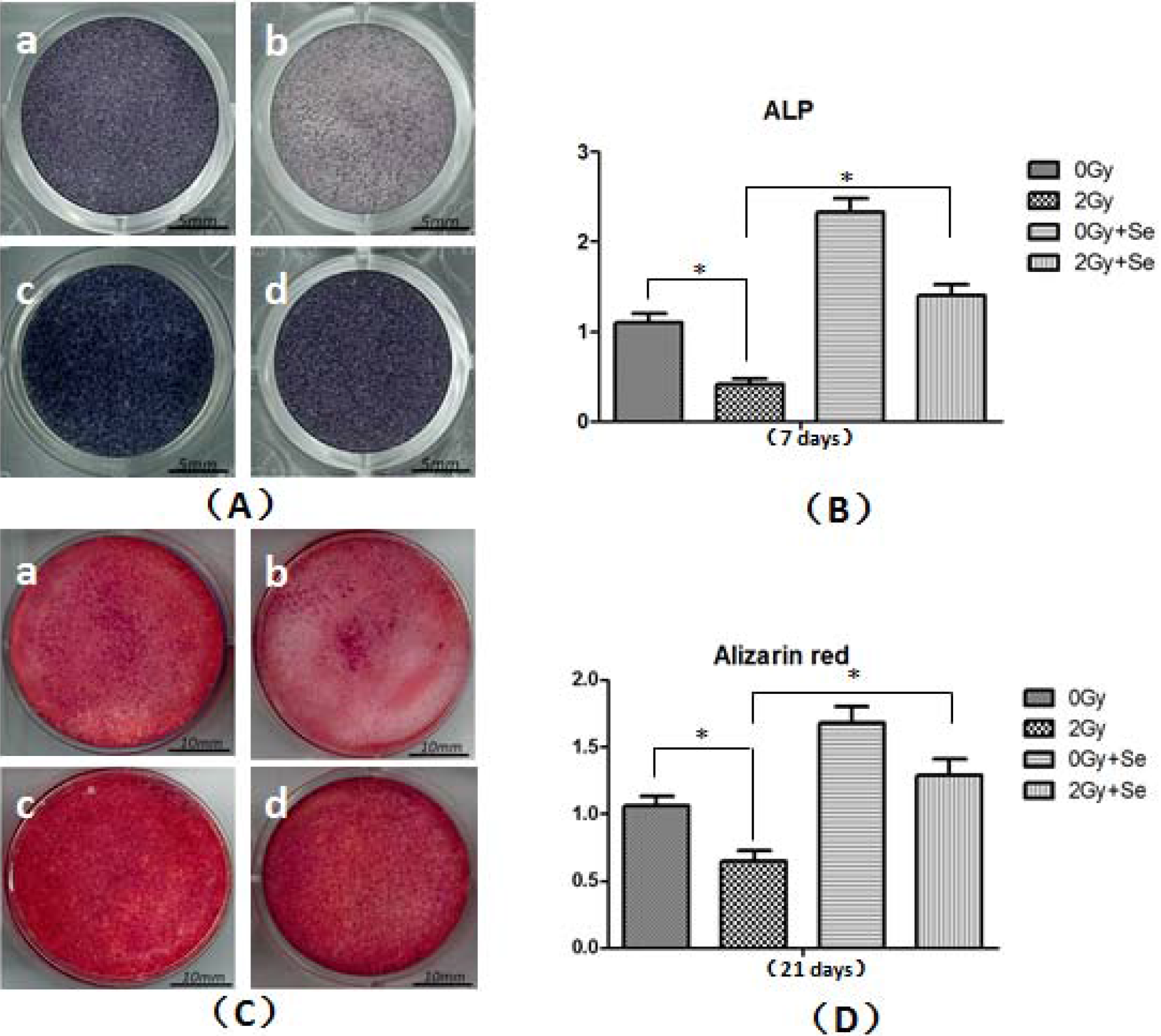
Effect of radiation and Sema3a on BMSC osteogenesis. (A) ALP activity was measured at 7 days after the corresponding treatment. The deeper the color, the stronger the activity. (B) Statistical data for ALP activity. (C) Effect of 2 Gy radiation and Sema3a on calcium deposition in BMSCs. (D) Statistical data for calcium deposition. a: 0 Gy; b: 2 Gy; c: 0 Gy + Sema3a; d: 2 Gy + Sema3a. *P < 0.05. All values are expressed as the mean ± SD, N = 5.

### 4.4 Effect of radiation and Sema3a on BMSC adipogenesis

At a dose of 2 Gy radiation, a greater number of Oil-red-O-positive cells were observed (Figure 4a). TG levels were increased by approximately 100% in group B compared with group A. However, the levels of TG were reduced by approximately 50% in group D compared with group B. The levels of TG were similar in groups A and D.

**Fig. 4.**
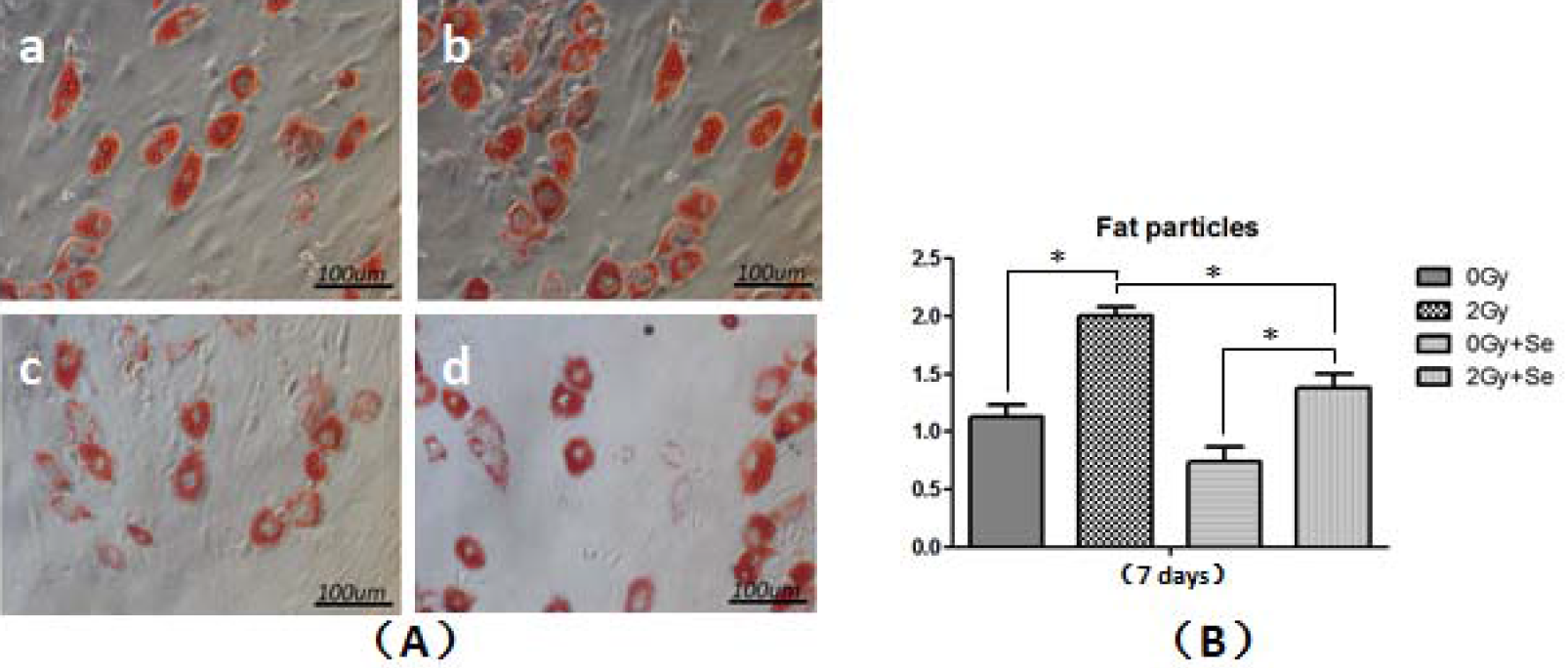
Effect of Sema3a and 2 Gy radiation on BMSC adipogenesis. (A) Effect of radiation and Sema3a on BMSC adipogenesis. (B) Statistical data for BMSC adipogenesis. a: 0 Gy; b: 2 Gy; c: 0 Gy + Sema3a; d: 2 Gy + Sema3a. *P < 0.05. All values are expressed as the mean ± SD, N = 5.

### 4.5 Effect of radiation and Sema3a on the cell cycle

The results of flow cytometry are depicted in Figure 5a, and the statistical data are shown in Figure 5b. The (S+G2) phase accounted for approximately 23%, 10%, 29%, and 20% of cells in groups A, B, C, and D, respectively. The results demonstrated that the number of cells in the division phase (S+G2) was lower in group B than in group A (P < 0.05); however, it was higher in group D than in group B (P < 0.05).

**Fig. 5.**
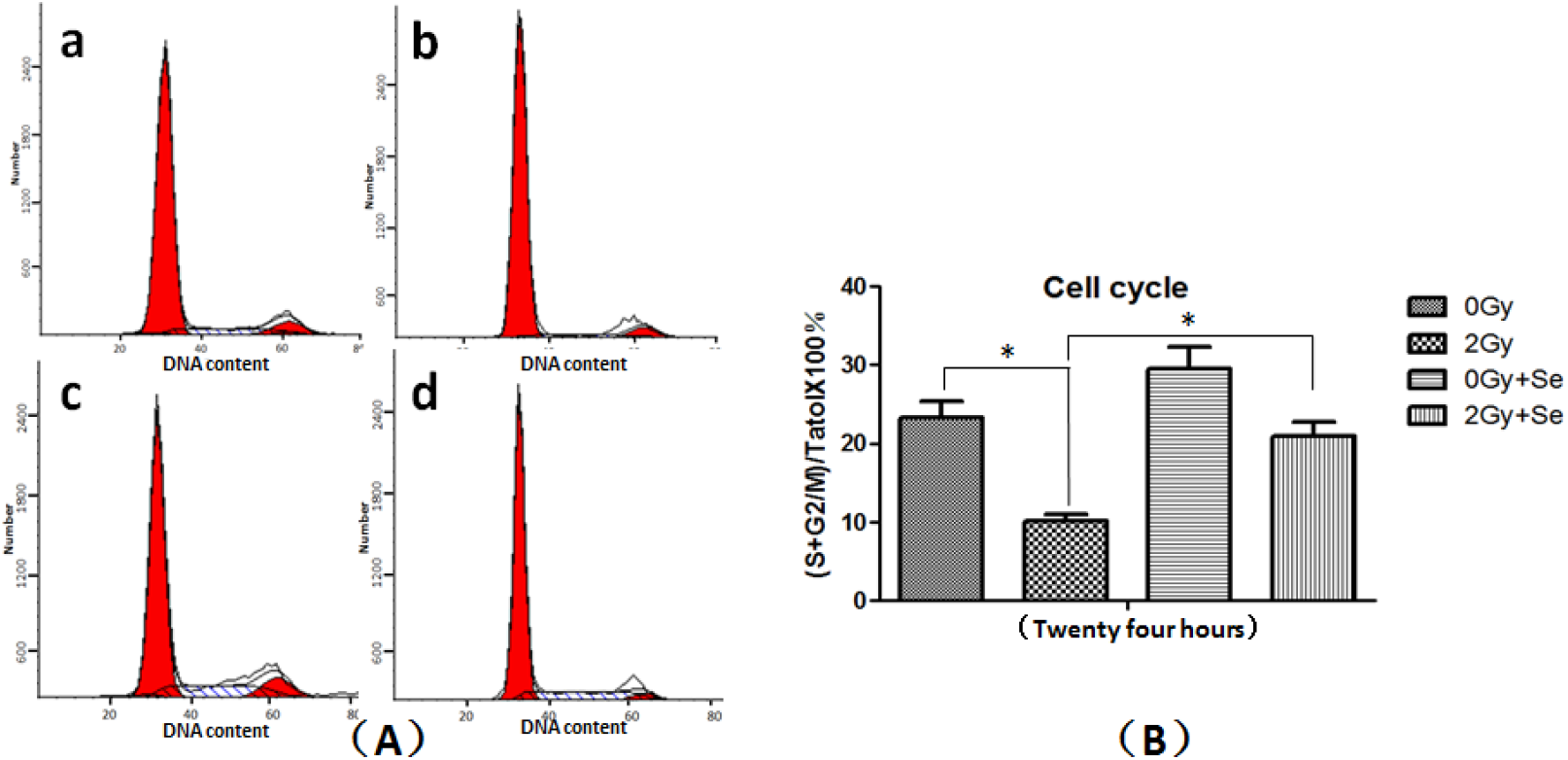
Effect of Sema3a and 2 Gy radiation on the cell cycle. (A) Effect of radiation and Sema3a on the cell cycle. (B) Statistical analysis of the cell cycle. a: 0 Gy; b: 2 Gy; C: 0 Gy + Sema3a; d: 2 Gy + Sema3a. *P < 0.05. All values are expressed as the mean ± SD, N = 5.

### 4.6 Effect of radiation and Sema3a on the generation of ROS in BMSCs

The DCFH-DA probe was used to measure the effect of 2 Gy radiation and Sema3a on the generation of ROS in BMSCs (Figure 6). At 2 h after exposure to radiation, a greater number of ROS-positive cells were observed in group B compared with the other groups, and the fluorescence of DCFH-DA, indicating ROS, was much brighter in group B than in the other three groups. However, the smallest number of ROS-positive cells and the lowest level of fluorescence were observed in group B. The data showed that the levels of ROS were two times higher in the 2 Gy group (group B) than in group A (P < 0.05). Additionally, the levels of ROS were much lower in group D than in group B (P < 0.05); Sema3a reduced the production of ROS by approximately 50% in group D compared with group B.

**Fig. 6.**
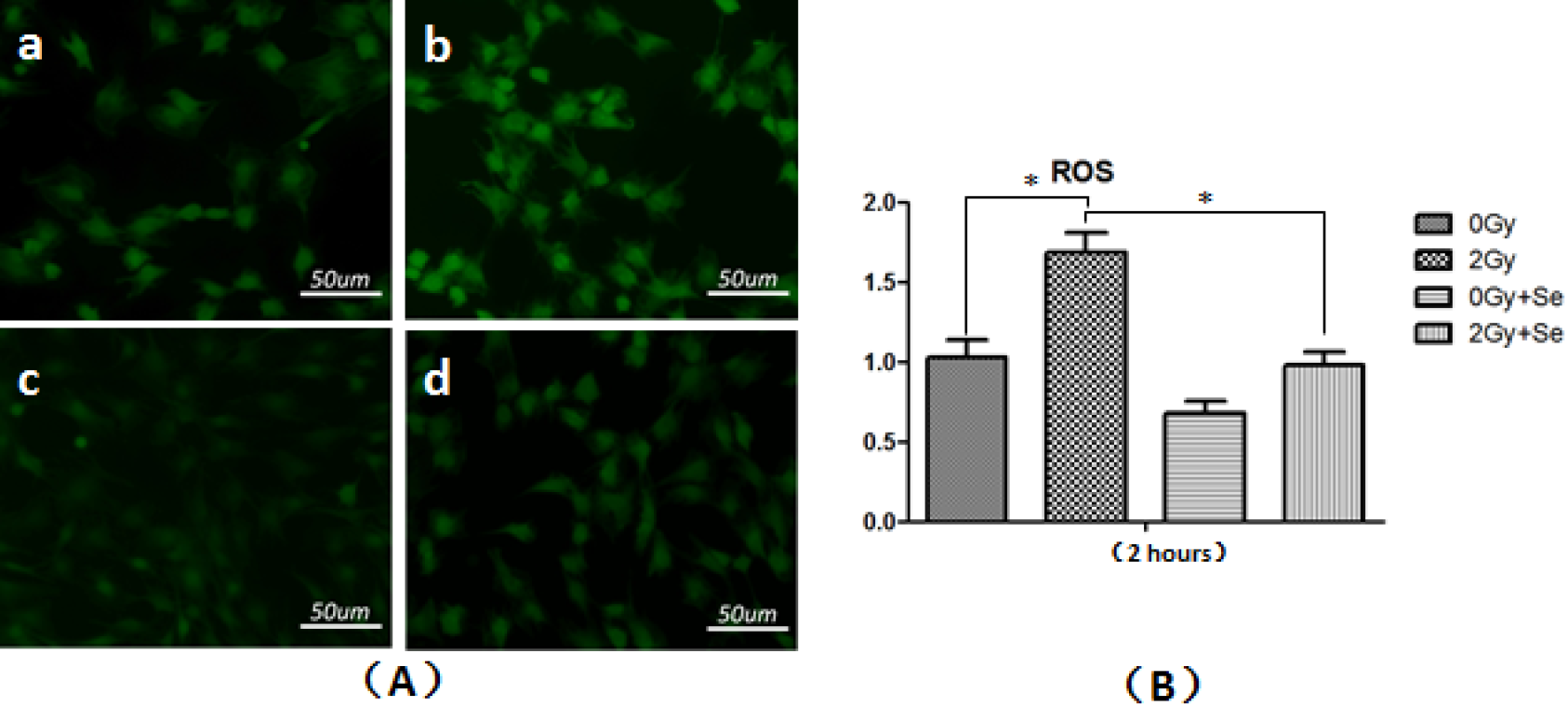
ROS generation in BMSCs. (A) Effect of radiation and Sema3a on ROS generation in BMSCs. Brighter green colors indicate higher levels of ROS. (B) Statistical data for ROS production in cells. a: 0 Gy; b: 2 Gy; c: 0 Gy + Sema3a; d: 2 Gy + Sema3a. *P < 0.05. All values are expressed as the mean ± SD, N = 5.

### 4.7 Effect of radiation and Sema3a on gene expression

The RNA levels of osteogenic (OCN and RUNX2) and adipogenic (PPARγ) genes on the 5th and 10th day post-irradiation are shown in Figure 7. RUNX2 expression was higher on the 5th day than on the 10th day, but OCN expression was higher on the 10th day than on the 5th day in the four groups. OCN and RUNX2 expression was lower in group B compared with group A both days (P < 0.05). On the 5th day, RUNX2 expression was higher in group D than in group B (P < 0.05). On the 10th day, OCN expression was lower in group B than in group D (P < 0.05). On the contrary, PPARγ expression was much higher in group B than in the other three groups on both days (P < 0.05); it was lower in group D than in group B on both days. TNF-α expression was higher on the 5th day than on the 10th day in groups B and D, but IL-6 expression was higher on the 5th day than on the 10th day. IL-6 and TNF-α expression was lower in group A than in group B on both days (P < 0.05); it was also higher in group B than in group D on both days (P < 0.05).

**Fig. 7.**
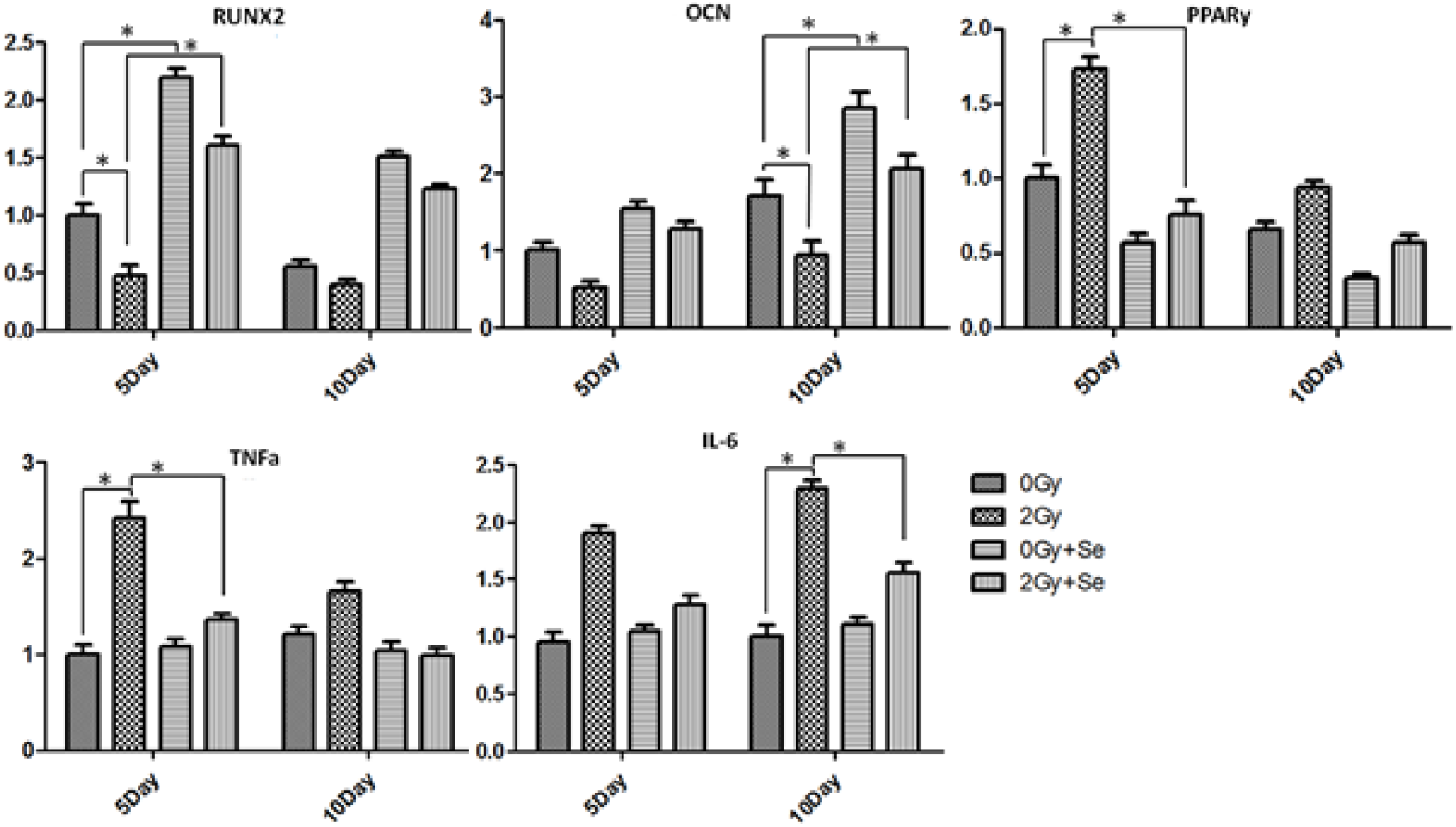
Expression of RUNX2, OCN, PPARγ, IL-6, and TNF-α on the 5th and 10th days. (A) RUNX2 expression. (B) OCN expression. (C) PPARγ expression. (D) TNF-α expression. (E) IL-6 expression. *P < 0.05. All values are expressed as the mean ± SD, N = 5.

## 5. Discussion/Conclusion

It is well known that radiotherapy leads to DNA damage in normal cells. If normal cells fail to repair the damaged DNA in time, they undergo apoptosis and cell cycle arrest [24–27]. In the clinical setting, radiotherapy is typically administered in fractions of 2 Gy. Many in vivo and in vitro studies generally use doses of 2 Gy for experimentation [25, 27]. The present study found that 2 Gy radiation could clearly inhibit cell proliferation, which was similar to the observations of previous studies [25–26]. Radiation-induced cell damage arises due to the energy deposited directly onto DNA and the induction of ROS, which cause DNA breakage in the nucleus and affect cell cycle checkpoints. Cell cycle checkpoints ensure the accuracy of DNA replication and division, and they also reflect the time needed to repair damaged DNA [28]. In addition, DNA damage is associated with many physiological processes of cells, including the activation of checkpoint kinase 1 (Chk1). Chk1 activation results in phosphorylation and the inactivation of cell division cycle 25 (cdc25), leading to the inactivation of the cdc2-B1 complex and S+G2/M arrest [29]. The results of the present study demonstrated that radiation administered at a dose of 2 Gy significantly increased the intracellular levels of ROS (Figure 7), while it significantly decreased the number of cells in the division (S+G2/M) phase of the cell cycle and significantly increased the number of cells in the quiescent stage (G1) (Figure 6). In order to repair DNA damage before cell division, the cell cycle DNA damage checkpoints occur late in the quiescent stage (G1), thereby preventing entry into the division phase (S+G2/M) [27,30]. Therefore, after cells receive radiation, the cell cycle arrests in the G1 phase.

Liu et al. demonstrated that Sema3a overexpression significantly promotes BMSC proliferation [31]. Although Sema3a showed a positive effect on cell proliferation in our study, the difference was not statistically significant; however, Sema3a significantly improved proliferation under 2 Gy radiation (Figure 2). Our study also demonstrated that the levels of ROS were significantly lower in group D than in group B (Figure 2). This means that Sema3a was able to reduce the production of ROS induced by radiation, thereby reducing the DNA damage caused by ROS and promoting cell proliferation. Analysis of the cell cycle checkpoints showed the same result (Figure 6). Zhao et al. reported that Sema3a deficiency could lead to cell apoptosis, the induction of hypoxia-induced myocardial injury by decreasing the secretion of hypoxia-induced inflammatory factors (TNF-α, IL-1β, and IL-6), the reduction of cell viability decline, and the inhibition of ROS production [32]. BMSCs are often used as seed cells due to their ability to differentiate into bone, cartilage, and adipose tissue. The osteogenic and adipogenic differentiation abilities of BMSCs are reciprocal [33]. BMSCs are sensitive to radiation in vivo and in vitro, and several studies have demonstrated that radiation decreases osteogenic differentiation, but increases adipogenic differentiation [34]. Our study showed that 2 Gy radiation significantly decreased ALP activity and calcium deposition compared with 0 Gy radiation. On the contrary, 2 Gy radiation increased the number of adipocyte islands and the levels of TG, indicating that 2 Gy radiation promoted the differentiation of BMSCs into adipocytes and inhibited their differentiation into osteoblasts. In vivo studies have shown that, in irradiated bone, there is more adipose tissue in bone marrow, and bone marrow is partly replaced by adipose tissue [35]. Furthermore, the expression of RUNX2 and OCN, which are bone formation-related genes, was much lower in the 2 Gy radiation group than in the 0 Gy group. Nevertheless, the expression of PPARγ, a lipogenesis-related gene, was much higher in the 2 Gy group than in the 0 Gy group. Some studies have also demonstrated that cells do not lose their differentiation ability completely when receiving high doses of radiation [33, 36]; however, radiation higher than 4 Gy significantly inhibits both adipogenic and osteogenic differentiation [33, 36].

Our research also demonstrated that Sema3a profoundly promoted osteogenic differentiation and inhibited adipogenic differentiation, which was similar to the findings of previous studies [17, 21, 37]. The canonical Wnt/β-catenin signaling pathway plays an important role in promoting osteogenesis and inhibiting adipogenesis. Sema3a activates Rac1, promoting β-catenin localization in the nucleus in response to Wnt ligands through FERM, RhoGEF, and FARP2, leading to the accumulation of β-catenin, which then activates the Wnt/β-catenin pathway [17, 21, 23, 37–38]. Previous studies indicated that radiation suppresses the Wnt/β-catenin pathway, inhibiting the differentiation of osteoblasts and reducing bone formation [34–35, 39–40]. In the present study, we found that Sema3a promoted ALP activity and calcium deposition, increased the expression of osteogenic-related genes, and decreased the formation of fat granules in group D compared with group B. It might be that Sema3a can reactivate the Wnt/β-catenin pathway, which was previously inhibited by radiation. At the same time, the expression of PPARγ, a key molecule for the promotion of adipogenic differentiation, was increased by radiation, but lowered when Sema3a was added.

Moreover, radiation increases the cellular levels of ROS and stimulates the production of inflammatory factors [21, 41–43]. Sema3a showed an inhibitory effect on ROS generation and the expression of inflammatory factors, which were induced by 2 Gy radiation. Previous studies have demonstrated that Sema3a acts as a potent immunosuppressive factor by regulating inflammatory responses. Sema3a, expressed by activated T cells and mesenchymal stem cells, inhibits T cell proliferation and cytokine secretion by binding to neuropilin-1 (NP-1), a neuronal receptor that is constitutively expressed on the surface of T cells and involved in the regulation of T cell proliferation through its main ligand Sema3a, thereby arresting T cells in the G0/G1 phase of the cell cycle [44–45]. Furthermore, Sema3a reduces the levels of anti-collagen IgG and suppresses the release of collagen-specific pro-inflammatory cytokines (interferon-γ and IL-17). However, it increases the expression of IL-10. The high concentration of IL-10 produced by Th2 cells suppresses the proliferation of Th1 cells in the serum [44, 46]. In the present study, we found that IL-6 and TNF-α expression was significantly decreased in group D compared with group B (Figure 7). Hence, Sema3a might reduce the release of inflammatory factors caused by radiation through the NP-1 signaling pathway and reduce the production of ROS to reduce the negative effects of radiation on BMSCs.

In addition, previous studies reported that Sema3a decreases osteoclastic cell differentiation and the activity of osteoclasts, and increases osteoclast apoptosis by inhibiting RANKL-induced tyrosine phosphorylation of phospholipase Cg2 and calcium oscillations through the immune-receptor tyrosine-based activation motif (ITAM) signaling pathway. RANK and ITAM signaling cooperates to induce the expression of NFATc1, a transcription factor for osteoclast-specific genes [47–48]. Additionally, Sema3a rescues bone loss in an ovariectomized mouse model of postmenopausal osteoporosis, increases callus volume and density at 4 weeks post-fracture, and promotes callus ossification and remodeling at 8 weeks post-fracture in osteoporotic rats [17, 21]. In vitro experiments have also shown that Sema3a suppresses osteoclastogenesis and promotes osteoblastogenesis [17, 21, 48–49]. Furthermore, Liu et al. demonstrated that Sema3a improves implant osseointegration and fixation in the proximal tibiae of ovariectomized rats [17]. These data suggest that Sema3a has the potential to be applied for the treatment of bone diseases caused by radiation.

## 6. Appendix

There is no appendix.

## 7. Supplementary Material

There is no supplementary material.

## 8. Statements

### 8.1. Acknowledgement

Thanks to the group of teachers in the radiology department of the Seventh People’s Hospital of Chengdu. This work was supported by National Natural Science Foundation of China (NO.813009061, NO.81500895, NO.81700941 and NO.81571008) and Sichuan Province Science and Technology Support Program (2016SZ0010).

### 8.2. Statement of Ethics

The animal experiment protocol was approved by the Animal Research Ethics Committee of West China Hospital of Stomatology, Sichuan University (NO.WCCSIRB-D-2015049)

### 8.5. Author Contributions

Ping Gong and Zheng Yang conceived the idea, designed and prepared samples. Bo Huang collected literature and recorded experimental data. Haiyang Tang proposed the experimental improvement programme. Haiyang Tang and Tao He analysed results and wrote the draft manuscript. Tao He provided overall guidance, supervised the experiments and revised the manuscript. All the authors gave their final approval for publication.

## References (Numerical)

[1] Hoskin PJ, Yarnold JR, Roos DR, et al (2001). Radiotherapy for bone metastases. Clin Oncol. 13:88–90.

[2] Shuja, Elghazaly, Iqbal, Mohamed R (2018).Efficacy of 8 Gy Single Fraction Palliative Radiatiion Therapy in Painful Bone Metastases: A SingleInstitution Experience. Cureus. 8:10(1).

[3] Jason A. Horton, Kathryn et al (2013). Mesenchymal Stem Cells Inhibit Cutaneous Radiation-Induced Fibrosis by Suppressing Chronic Inflammation. Stem cells. 31:2231–2241.

[4] Bo Gong, Megan E. Oest, Kenneth A. Mann et al (2013). Raman Spectroscopy Demonstrates Prolonged Alteration of Bone Chemical Composition Following Extremity Localized Irradiation. Bone. 57(1): 252–258.

[5] Priestman TJ, Bullimore JA, Godden TP, Deutsch GP (1989). The Royal College of Radiologists’ Fractionation Survey. Clin Oncol.1:39–46.

[6] S. Chauhan, S. A. Khan.A, Prasad (2018). Irradiation-Induced Compositional Effects on Human Bone After Extracorporeal Therapy for Bone Sarcoma. Calcif TissueInt. doi:10.1007/s00223-018-0408-2.

[7] Green DE, Adler BJ, Chan ME, Rubin CT (2012). Devastation of adult stem cell pools by irradiation precedes collapse of trabecular bone quality and quantity. J Bone Miner Res. 27:749–59.

[8] Willey JS, Livingston EW, Robbins ME, Bourland JD, Tirado-Lee L, Smith-Sielicki H, Bateman TA (2010). Risedronate prevents early radiation-induced osteoporosis in mice at multiple skeletal locations. Bone. 46(1): 101 – 111.

[9] Kelly J, Damron T, Grant W, Anker C, Holdridge S, Shaw S, Horton J, Cherrick I, Spadaro J (2005).Cross-sectional study of bone mineral density in adult survivors of solid pediatric cancers. J Pediatr Hematol Oncol. 27:248–53.

[10] Jeffrey S. Willey, Shane A. J. Lloyd, Gregory A. Nelson, Ted A. Batem (2011); Ionizing Radiation and Bone Loss: Space Exploration and Clinical Therapy Applications; Clinic Rev Bone Miner Meta b. 9:54–62.

[11] Helmstedter CS, Goebel M, Zlotecki R, Scarborough MT (2011). Pathologic fractures after surgery and radiation for soft tissue tumors. Clin Orthop Relat Res. 1:165–172.

[12] Bassam Shugaa-Addin, Hashem-Motahir Al-Shamiri (2016); The effect of radiotherapy on survival of dental implants in head and neck cancer patients. J ClinExp Dent. 8(2):e194–200.

[13] Gortzak Y, Lockwood GA, Mahendra A, Wang Y, Chung PW, Catton CN, O’Sullivan B, Deheshi BM, Wunder JS, Ferguson PC (2010). Prediction of pathologic fracture risk of the femur after combined modality treatment of soft tissue sarcoma of the thigh. Cancer. 116:1553–9.

[14] Brasseur M, Brogniez V, Grégoire V, Reychler H, Lengelé B, D’Hoore W, et al (2010) Effects of irradiation on bone remodeling around mandibular implants: an experimental study in dogs. Int J Oral Maxillofac Surg. 35:850–855.

[15] Divya Devaraj, d. Srisakthi (2014); Hyperbaric Oxygen Therapy – Can It Be the New Era in Dentistry?; Journal of Clinical and Diagnostic Research. Vol-8(2):263–265.

[16] Fukuda T, Takeda S, Xu R, Ochi H, Sunamura S, Sato T et al (2013). Sema3a regulates bone-mass accrual through sensory innervations. Nature.497:490–493.

[17] Hayashi M, Nakashima T, Taniguchi M, Kodama T, Kumanogoh A, Takayanagi (2012). Osteoprotection by semaphorin 3A. Nature. 485:69–74.

[18] Verlinden L, Kriebitzsch C, Beullens I, Tan BK, Carmeliet G, Verstuyf A (2013). Nrp2 deficiency leads to trabecular bone loss and is accompanied by enhanced osteoclast and reduced osteoblast numbers. Bone. 55:465–475.

[19] Sujin Kanga, Atsushi Kumanogoha (2014). Semaphorins in bone development, homeostasis, and disease. Seminars in Cell & Developmental Biology. 24: 163– 171.

[20] Natalie A Sims and T John Martin (2014); Coupling the activities of bone formation and resorption: a multitude of signals within the basic multicellular unit. BoneKEy Reports. 3: 481.

[21] Ren Xu (2014); Semaphorin 3A: A new player in bone remodeling; Cell Adhesion & Migration. 8:1, 5–10.

[22] Yunfeng Li, Jing Hu et al (2015). The effect of semaphorin 3A on fracture healing in osteoporotic rats. Journal of Orthopaedic Science. 20 (6): H14–1121.

[23] Zhenxia Li, Jin Hao, Xin Duan, Nan Wu, Zongke Zhou et al (2017), The Role of Semaphorin 3A in Bone Remodeling. Frontiers in Cellular Neuroscience. 2:40.

[24] C. Ferlini, R. D’Amelio, G. Scambia (2012). Apoptosis induced by ionizing radiation. The biological basis of radiosensitivity. SubCellular Biochemistry, vol. 36:171–186.

[25] W. Su, Y. Chen, W. Zeng, W. Liu, H. Sun (2012). Involvement of Wnt signaling in the injury of murine mesenchymal stem cells exposed to X-radiation. International Journal of Radiation Biology, vol. 88, no. 9, pp. 635–641.

[26] Yuan Y, Yan G, Gong R, Zhang L, Liu T, Feng C et al (2017). Effects of Blue Light Emitting Diode Irradiation On the Proliferation, Apoptosis and Differentiation of Bone Marrow-Derived Mesenchymal Stem Cells. Cell Physiol Biochem. 43(1):237–246. doi: 10.1159/000480344.

[27] Amir Sabet Sarvestani, Parviz Abdolmaleki, Seyed Javad Mowla et. al (2010). Static magnetic fields aggravate the effects of ionizing radiation on cell cycle progression in bone marrow stem cells. Micron. 41:101–104.

[28] Sherr C.J (2000). The Pezcoller lecture: cancer cell-cycles revisited. Cancer Res. 60, 3695–3698.

[29] Jackson J.R, Gilmartin, A, Imburgia, C, Winkler, J.D, Marshall, L.A., Roshak, A (2010). An indolocarbazole inhibitor of human checkpoint kinase (Chk1) abrogates cell cycle arrest caused by DNA damage. Cancer Res. Feb 1;60(3):566–72.

[30] Ji K, Sun X, Liu Y, Du L, Wang Y, He N et al (2018). Regulation of Apoptosis and Radiation Sensitization in Lung Cancer Cells via the Sirt1/NF-κB/Smac Pathway. Cell Physiol Biochem. Jul 17;48(1):304–316. doi: 10.1159/000491730.

[31] Li Liu, Jue Wang, Xiaomeng Song (2018). Semaphorin 3A promotes osteogenic differentiation in human alveolar bone marrow mesenchymal stem cells. Experimental and therapeutic medicine. 15: 3489–3494.

[32] Zhao C, Liu J, Zhang M, Wu Y (2016). Semaphorin 3A deficiency improves hypoxia-induced myocardial injury via resisting inflammation and cardiomyocytes apoptosis. Cell Mol Biol. Feb 4;62(2):8–l4.

[33] J. N. Beresford, J. H. Bennett, C. Devlin, P. S. Leboy, M. E. Owen (1992). Evidence for an inverse relationship between the differentiation of adipocytic and osteogenic cells in rat marrow stromal cell cultures. Journal of Cell Science, vol. 102, no. 2, pp,341–351.

[34] F. Mussano, K. J. Lee, P. Zuk et al (2010). Differential effect of ionizing radiation exposure on multipotent and differentiation restricted bone marrow mesenchymal stem cells. Journal of Cellular Biochemistry, vol. 111, no. 2, pp. 322–332.

[35] X. Cao, X. Wu, D. Frassica et al (2011). Irradiation induces bone injury by damaging bone marrow microenvironment for stem cells. Proceedings of the National Academy of Sciences of the United States of America, vol. 108, no. 4, pp. 1609–1614.

[36] M.-F. Chen, C.-T. Lin, W.-C. Chen et al (2006). The sensitivity of human mesenchymal stem cells to ionizing radiation. International Journal of Radiation Oncology Biology Physics, vol. 66, no. 1, pp. 244–253.

[37] Tang P, Yin P, Lv H, Zhang L, Zhang L (2015). The Role Of Semaphorin 3A In The Skeletal System. Crit Rev Eukaryot Gene Expr. 25(1):47–57.

[38] R. Baron and M. Kneissel (2013). WNT signaling in bone homeostasis and disease: from human mutations to treatments. Nature Medidne. vol. 19, no. 2, pp. 179–192.

[39] Xiangwei Liu, Naiwen Tan, Yuchao Zhou et al (2016). Semaphorin 3A Shifts Adipose Mesenchymal Stem Cells towards Osteogenic Phenotype and Promotes Bone Regeneration In Vivo. Stem Cells International. Volume, http://dx.doi.org/10.1155/2016/2545214.

[40] Lee SY, Jeong EK, Ju MK, Jeon HM, Kim MY, Kim CH, Park HG, Han SI, Kang HS (2017). Induction of metastasis, cancer stem cell phenotype, and oncogenic metabolism in cancer cells by ionizing radiation. Mol Cancer. Jan 30; 16(1): 10. doi: 10.1186/s12943-016-0577-4.

[41] Muzaffer U, Paul VI, Prasad NR, Karthikeyan R, Agilan B, Protective effect of Juglans regia L (2018). against ultraviolet B radiation induced inflammatory responses in human epidermal keratinocytes. Phytomedicine. Mar 15;42:100–111.

[42] Debayan Mukherjee, Philip J Coates, Sally A Lorimore et al (2014). Responses to ionizing radiation mediated by inflammatory mechanisms; J Pathol. 232: 289–299.

[43] Riley, P.A (1994). Free radicals in biology: oxidative stress sand the effects of ionizing radiation. Int. J.Radiat.Biol. 65:27–33.

[44] Yamamoto, M, K. Suzuki, T. Okuno, T. Ogata, N. Takegahara, H. Takamatsu, M. Mizui, M. Taniguchi, A. Che’dotal, F. Suto, et al (2008). Plexin-A4 negatively regulates T lymphocyte responses. Int. Immunol. 20: 413–420.

[45] Lepelletier Y, Lecourt S, Renand A, Arnulf B, et.al (2010).Galectin-1 and semaphorin-3A are two soluble factors conferring T-cell immunosuppression to bone marrow mes nchymal stem cell. Stem Cells Dev. Jul; 19(7): 1075–9. doi: 10.1089/scd.2009.0212.

[46] Catalano, A., P. Caprari, S. Moretti, M. Faronato, L. Tamagnone, and A. Procopio (2006). Semaphorin-3A is expressed by tumor cells and alters T-cell signal transduction and function. Blood. 107: 3321–3329.

[47] Yamashita, T, Takahashi, N, and Udagawa, N (2012). New roles of osteoblasts involved in osteoclast differentiation. World J. Orthop. 3:175–181. doi: 10.5312/wjo.v3.i11.175.

[48] Li, Y. Yang, L., He, S., and Hu, J (2015). The effect of semaphorin 3A on fracture healing in osteoporotic rats. J. Orthop. Sci. 20, 1114–1121. doi: 10.1007/s00776-015-0771-z.

[49] Li Y, He D, Liu B, Hu J (2017).SEMA3A suspended in matrigel improves titanium implant fixation inovariectomized rats. J Biomed Mater Res B Appl Biomater. Oct, 105(7):2060–2065. doi:10.1002/jbm.b.33730.

[50] Santivasi, W.L., Xia, F (2014). Ionizing radiation-induced DNA damage, response, and repair. Antioxid.Redox Signal. 21 (July 2), 251–259.

